# Altered motoneuron properties contribute to motor deficits in a rabbit hypoxia ischemia model of cerebral palsy

**DOI:** 10.1101/817957

**Authors:** P. Steele, C. F. Cavarsan, L. Dowaliby, M. Westefeld, A. Drobyshevsky, M. A. Gorassini, K. A. Quinlan

**Author notes:** Corresponding Author: Katharina A. Quinlan, 130 Flagg Rd, Kingston, RI 02881.

## Abstract

Cerebral palsy (CP) is caused by a variety of factors attributed to early brain damage, resulting in permanently impaired motor control, marked by weakness and muscle stiffness. To find out if altered physiology of spinal motoneurons (MNs) could contribute to movement deficits, we performed whole cell patch clamp in neonatal rabbit spinal cord slices after developmental injury at 79% gestation. After preterm hypoxia-ischemia (HI), rabbits are born with motor deficits consistent with a spastic phenotype including hypertonia and hyperreflexia. There is a range in severity, thus kits are classified as severely affected, mildly affected, or unaffected based on modified Ashworth scores and other behavioral tests. At postnatal day (P)0-5, we recorded electrophysiological parameters of 40 MNs in transverse spinal cord slices using whole cell patch clamp. Using a multivariate analysis of neuronal parameters, we found significant differences between groups (severe, mild, unaffected and sham control MNs), age (P0 to P5) and spinal cord region (cervical to sacral). Severe HI MNs showed more sustained firing patterns, depolarized resting membrane potential, and a higher threshold for action potentials. These properties could contribute to both muscle stiffness and weakness, respectively, hallmarks of spastic CP. Interestingly altered persistent inward currents (PICs) and morphology in severe HI MNs would dampen excitability (reduced normalized PIC amplitude and increased dendritic length). In summary, changes we observed in spinal MN physiology likely contribute to severity of the phenotype including weakness and hypertonia, and therapeutic strategies for CP could target excitability of spinal MNs.

**Key Points:**

- Physiology of neonatal spinal motoneurons is altered after *in utero* hypoxia-ischemic injury
- In motoneurons from severely affected animals there is more sustained firing (lower ΔI values), a depolarized resting potential, but a higher voltage threshold for action potential firing.
- Altered motoneuron excitability could contribute directly to muscle stiffness and spasticity in cerebral palsy.

## Introduction

Cerebral palsy is not well understood, despite its prevalence and seriousness. There exist only a few evidence-based treatments for cerebral palsy: the effectiveness of many currently used therapeutic strategies is unclear (46, 58). Recent clinical advances include use of magnesium sulfate and hypothermia after hypoxic-ischemic encephalopathy to acutely reduce neural damage (41, 51, 55, 60), but little basic research is devoted to addressing symptoms after they arise. Part of the problem in treating CP may be the diversity of causes including neonatal stroke, placental insufficiency, preterm birth, inflammation, traumatic injury, difficulties during birth and many other contributing factors (25, 40). Another problem could be that modeling the condition in animals is complicated, and while rodent models are useful for development of neuroprotective strategies, larger animal models are needed to study motor deficits (9, 11).

Loss of corticospinal control of movement is considered causative of motor deficits in CP, but little investigation into the precise effect on spinal circuits has been conducted. A notable exception is the work of John H. Martin and colleagues, who have documented changes in corticospinal synaptic connectivity in specific spinal laminae and loss of cholinergic interneurons after cortical silencing during development or lesions of the corticospinal tract (20–22, 34, 35, 37, 43). Another important study showed changes in parvalbumin-positive spinal interneurons after cortical silencing in development (10, 12). Both of these interneuron classes are synaptically connected to spinal MNs, and could contribute to altered motor output. Based on these foundational studies, our hypothesis was that altering development with HI injury would also alter development of MNs, specifically the electrophysiological properties governing excitability in spinal MNs. We further hypothesized that changes in excitability would correspond / contribute to the severity of motor deficits. In short, that altered activity of spinal MNs could contribute to muscle stiffness and spasticity.

In order to assess changes in intrinsic properties of spinal MNs, we used the rabbit HI model of cerebral palsy (14). It’s been shown in previous studies that HI injury during late gestation in rabbits can result in a variety of neurologic and muscular damage, including muscle stiffness (14), loss of neurons in cortical layers 3 and 5, white matter injury, thinning of the corticospinal tract (8), cell death in the spinal cord and decreased numbers of spinal MNs (15), increased sarcomere length, decreased muscle mass and hyperreflexia (54). There is also an increase in spinal monoamines which could increase excitability of spinal neurons and thus promote spasticity (3, 16). Thus, changes observed in spinal MNs in the rabbit model could be directly compared to motor deficits.

Changes in MN physiology are likely to contribute to motor impairment in cerebral palsy, yet this has not been directly assessed in any animal models. Thus, we assessed electrophysiological parameters in spinal MNs in neonatal rabbits after sham surgery or hypoxic-ischemic insult during development.

## Methods

All rabbits were used according to both the University of Rhode Island’s, Northwestern University’s and Northshore University Health System’s Animal Care and Use Committee guidelines. Pregnant New Zealand White rabbits (Charles River Laboratories, Inc, Wilmington MA), underwent HI procedures as described in (14). Briefly, at ~80% gestation (day 25 of gestation (E25) dams were anesthetized, and the left femoral artery was isolated. A Fogarty balloon catheter was inserted into the femoral and advanced to the level of the descending aorta, above the uterine arteries and inflated for 40 minutes. Sham animals underwent the same procedures but without inflation of the catheter. After the procedure, the dam recovered and later gave birth to kits with HI injuries. Categorization of the severity of the phenotype was performed by a blinded observer, using a modified Ashworth scale, observation / tests for activity, locomotion, posture, righting reflex, muscle tone (as described in (14)). One rabbit kit which was affected by HI but displayed a phenotype of hypotonia instead of hypertonia was removed from the data set. All other kits included in this study displayed hypertonic phenotype if affected by HI.

### Patch Clamp

Whole cell patch clamp was performed similar to previously published work (50) from P0-5. Briefly, horizontal spinal cord slices 350μm thick were obtained using a Leica 1000 vibratome. Slices were incubated for one hour at 30°C and perfused with oxygenated (95% O_2_ and 5% CO_2_) modified Ringer’s solution containing (in mM): 111 NaCl, 3.09 KCl, 25.0 NaHCO3, 1.10 KH2PO4, 1.26 MgSO4, 2.52 CaCl2, and 11.1 glucose at 2 ml/min. Whole cell patch electrodes (1-3 MΩ) contained (in mM) 138 K-gluconate, 10 HEPES, 5 ATP-Mg, 0.3 GTP-Li and Texas Red dextran (150μM, 3000MW). PICs were measured in voltage clamp mode with holding potential of −90mV and depolarizing voltage ramps of both 36 mV/s and 11.25 mV/s bringing the cell to 0mV in 2.5s or 8s, respectively and then back to the holding potential in the following 2.5s or 8s. Input resistance was measured from the slope of the leak current near the holding potential. Capacitance was measured with Multiclamp’s whole cell capacitance compensation function. Resting membrane potential was measured in voltage clamp as the voltage at which there is 0pA of injected current in the descending ramp. In current clamp, frequency – current measurements were obtained from current ramps. The first spike on the current ramp was used to measure properties of action potentials. Threshold voltage was defined as the voltage at which the action potential slope exceeds 10V/s. Rate of rise and fall of the action potential were measured as peak and trough of the first derivative of the action potential. Duration of the action potential was measured at half-peak (defined as the midpoint between overshoot and threshold voltages). Depolarizing current steps of varying amplitude were used to find maximum firing rates (near depolarization block) and to measure after-spike after hyperpolarization (in single spikes elicited near threshold). Hyperpolarizing current steps (typically between −850 and −1250pA) were used to measure hyperpolization-activated sag currents (IH). Neuron selection: Neurons were targeted in MN pools mainly from cervical and lumbar regions of the cord and were removed from the data set if their resting membrane potential was more depolarized than −45mV in current clamp.

### Imaging

After electrophysiological measurements were obtained, MNs were imaged to assess anatomical development, and photos were obtained of the electrode placement within the spinal cord slice, as shown in **Figure 1**. Images were acquired with a Nikon microscope fitted with a 40x water-dipping objective lens and two photon excitation fluorescence microscopy performed with a galvanometer-based Coherent Chameleon Ultra II laser. To optimize excitation of red/green fluorophores, the laser was tuned to 900nm. 3D reconstructions of MNs were created using Neurolucida 360° software.

**Figure 1.**
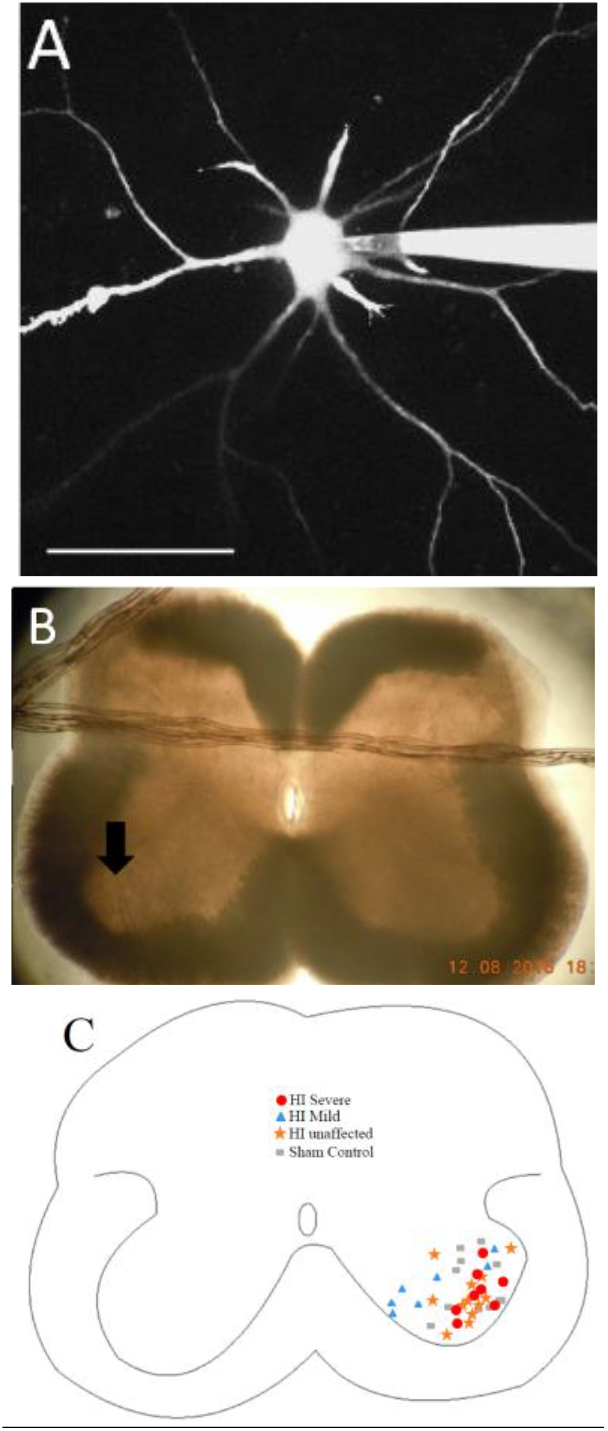
Patch clamp of MNs with dye-filling via patch electrode as in shown in **(A)**. Scale bar = 100μm. **(B)** Placement of patch electrode (at arrow) within the slice is captured with a photo. **(C)** Map of recorded MNs within medial and lateral motor pools.

### Statistics

A multivariate analysis using SPSS software was used for determining significance of parameters over groups. Injury classification (sham/control, HI unaffected, HI mildly affected, and HI severely affected), age of the kit (P0 – P5), and spinal cord region (cervical, thoracic, lumbar or sacral) were all included as fixed factors. Significance was determined by *p* values < 0.05.

## Results

After HI surgery was performed in pregnant dams at 79% gestation, kits were born naturally about a week later. At ages P0-5, neonates were tested for deficits based on modified Ashworth scores for stiffness in the limbs, postural deficits, and righting reflex. Kits could be given a maximum score of 6 (normal posture, righting and joint resistance). Since there was a large variation in the severity of motor deficits, HI kits were divided into 3 groups: HI unaffected (scores were the same range as control kits, 5-6), HI mild (scores 3-4), HI severe (scores 1-2). Experiments were all performed in the first 5 days of life. Over 40 spinal MNs were patched in transverse spinal cord slices, and over 40 parameters were measured from each. To determine significance of the variables, a multivariate analysis was used that included 3 independent variables: 1) condition (sham control, HI unaffected, HI mildly affected, and HI severely affected, 2) age (postnatal day 0-5), and 3) spinal region (cervical through sacral). All data, including mean, standard deviation, group size and *p* value is included in table format (Tables 1 – 16).

### HI MNs show sustained firing but higher voltage threshold

In rabbit kits severely injured by HI, MNs had significantly increased sustained firing. The frequency current (F-I) relationship was measured using current ramps, as shown in **Figure 2**. Depolarizing current ramps are used to evoke firing, and current at onset and offset of firing (I_ON_ and I_OFF_) determine ΔI. In sham control MNs, ΔI was larger and always a positive value (219pA mean, standard deviation 224pA), indicating firing ceased at a higher current amplitude on the descending ramp than the current level that elicited firing on the ascending ramp (see figure 2A). Severe HI MNs had a smaller, and usually negative ΔI (−16pA mean, standard deviation 84pA), revealing increasingly sustained firing (see figure 2B).

**Figure 2:**
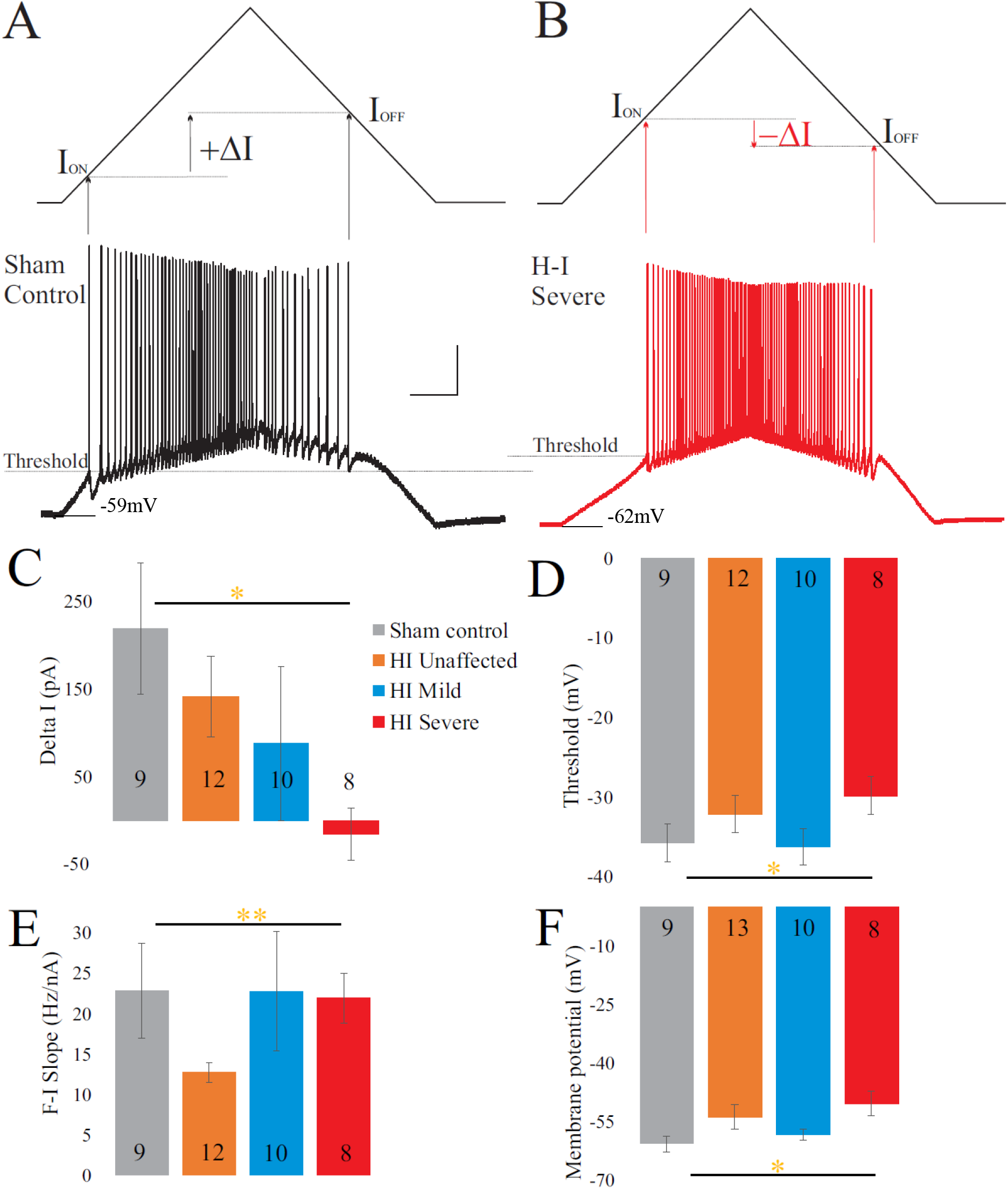
Severe HI MNs show more sustained firing than sham control MNs, but a higher voltage threshold. Control motoneurons **(A)** have larger values for ΔI compared to severe HI **(B)**. Average ΔI and threshold are shown in **(C)** and **(D)** for all groups. There was a significant group effect on slope of the frequency current relationship **(E)** indicating this is altered in HI unaffected MNs. Resting membrane potential was significantly more depolarized in HI severe MNs than sham control MNs **(F)**. Error bars = SEM. Scale bars in A = 20mV (vertical) and 0.5s (horizontal) and applies to panels A and B.

Threshold for action potential initiation occurred at a higher threshold (Figure 2A, B and D). Thus, MNs from severely affected kits could not be classified as hyperexcitable, since they required more depolarization to begin firing. However resting membrane potential was significantly more depolarized in HI MNs than sham controls. Mean values for these parameters are shown in Figure 2C, D, and F along with the intermediate groups (HI unaffected and HI mildly affected). While the most dramatic changes in MN physiology took place in the severely affected animals, there were interesting differences in “unaffected” and mildly affected neonates as well. The slope of the frequency – current relationship was smaller in unaffected MNs (see Fig 2E), indicating less rate modulation would be possible in these kits. Values for voltage threshold and membrane potential were also moderately depolarized in unaffected MNs (compare Fig 2D and 2F). In summary, HI during fetal development had an impact on MN physiology. MNs from rabbit kits that had severe phenotype showed the most dramatic changes, including the most depolarized resting membrane potential of all the groups (−50.5mV compared to −60.8mV mean in sham MNs), most sustained firing (219pA mean ΔI in sham controls vs −16pA in HI severe), and largest change in capacitance (mean 192pF in sham controls vs 282pF mean in HI severe), see Tables 1–4 for complete data sets.

**Table 1.** Frequency – current characteristics

| **Variable** | **Condition** | **Mean** | **SD** | **N** |
| --- | --- | --- | --- | --- |
| **I<sub>ON</sub> (pA)** | sham control | 213 | 153 | 9 |
|  | HI unaffected | 627 | 475 | 12 |
|  | HI mild | 422 | 458 | 10 |
|  | HI severe | 459 | 375 | 8 |
| **I<sub>OFF</sub> (pA)** | sham control | 416 | 357 | 9 |
|  | HI unaffected | 768 | 538 | 12 |
|  | HI mild | 509 | 319 | 10 |
|  | HI severe | 443 | 452 | 8 |
| **Δ I (pA)** | sham control | 219 | 224 | 9 |
|  | HI unaffected | 141 | 159 | 12 |
|  | HI mild | 88 | 279 | 10 |
|  | HI severe | -16 | 84 | 8 |
| **FI Slope (Hz/nA)** | sham control | 23 | 18 | 9 |
|  | HI unaffected | 13 | 4 | 12 |
|  | HI mild | 23 | 23 | 10 |
|  | HI severe | 20 | 9 | 8 |
| **Threshold (mV)** | sham control | -35.8 | 7.2 | 9 |
|  | HI unaffected | -32.2 | 8.0 | 12 |
|  | HI mild | -36.3 | 7.1 | 10 |
|  | HI severe | -29.9 | 6.7 | 8 |
| **Maximum instantaneous firing frequency (Hz)** | sham control | 98 | 28 | 9 |
|  | HI unaffected | 106 | 32 | 12 |
|  | HI mild | 129 | 33 | 10 |
|  | HI severe | 129 | 23 | 8 |
| **Maximum steady state firing frequency (Hz)** | sham control | 30 | 10 | 9 |
|  | HI unaffected | 36 | 13 | 12 |
|  | HI mild | 45 | 18 | 10 |
|  | HI severe | 37 | 13 | 8 |

**Table 2.** Significance of frequency – current variables

| **Fixed Factor** | **Variable** | **p** |
| --- | --- | --- |
| Condition<br>(Sham control, HI<br>unaffected, HI<br>mildly affected, HI<br>severely affected) | ION | .145 |
|  | IOFF | .225 |
| | $\Delta I$ | .010 |
|  | SLOPE | .007 |
|  | Threshold | .046 |
|  | Instantaneous firing freq | .176 |
|  | Steady state firing freq | .155 |
| AGE<br>(P0-5) | ION | .278 |
|  | IOFF | .413 |
| | $\Delta I$ | .020 |
|  | SLOPE | .937 |
|  | Threshold | .465 |
|  | Instantaneous firing freq | .105 |
|  | Steady state firing freq | .575 |
| Region<br>(cervical, thoracic,<br>lumbar, sacral) | ION | .676 |
|  | IOFF | .421 |
| | $\Delta I$ | .157 |
|  | SLOPE | .333 |
|  | Threshold | .820 |
|  | Instantaneous firing freq | .009 |
|  | Steady state firing freq | .435 |

**Table 3.** Intrinsic neuron properties

| *Variable* | *Condition* | *Mean* | *SD* | *N* |
| --- | --- | --- | --- | --- |
| Resting membrane potential (mV) | sham control | -60.8 | 6.4 | 9 |
|  | HI unaffected | -53.9 | 11.3 | 13 |
|  | HI mild | -58.5 | 4.3 | 10 |
|  | HI severe | -50.5 | 9.1 | 8 |
| Input resistance (MΩ) | sham control | 61.8 | 43.8 | 9 |
|  | HI unaffected | 34.9 | 15.6 | 13 |
|  | HI mild | 58.4 | 41.7 | 10 |
|  | HI severe | 43.8 | 13.4 | 8 |
| Capacitance (pF) | sham control | 192 | 98 | 9 |
|  | HI unaffected | 259 | 45 | 13 |
|  | HI mild | 214 | 77 | 10 |
|  | HI severe | 282 | 0 | 8 |

**Table 4.**
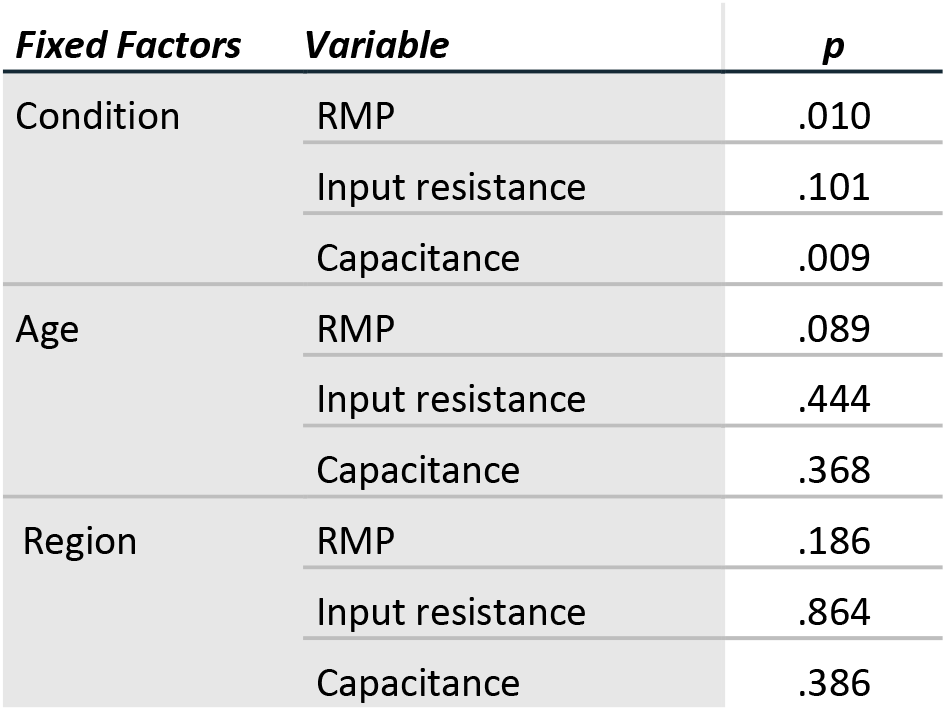
Significance of intrinsic neuron properties

| *Fixed Factors* | *Variable* | *p* |
| --- | --- | --- |
| Condition | RMP | .010 |
|  | Input resistance | .101 |
|  | Capacitance | .009 |
| Age | RMP | .089 |
|  | Input resistance | .444 |
|  | Capacitance | .368 |
| Region | RMP | .186 |
|  | Input resistance | .864 |
|  | Capacitance | .386 |

### Other changes in active properties were not present

A complete analysis of I_H_ (sag and rebound currents), action potential parameters, and after-spike after hyperpolarization (AHP) was performed and no significant differences in these parameters were found between groups. All data is included in Tables 9 – 12.

**Table 5.** Persistent inward current characteristics (5s ramp)

| **Variable** | **Condition** | **Mean** | **SD** | **N** |
| --- | --- | --- | --- | --- |
| Norm PIC amp<br>(pA/pF) | Sham | -2.10 | 2.44 | 9 |
|  | HI Unaffected | -1.45 | 0.81 | 13 |
|  | HI Mild | -1.86 | 1.02 | 9 |
|  | HI Severe | -1.22 | 0.41 | 8 |
| PIC onset (mV) | Sham | -42.9 | 7.3 | 9 |
|  | HI Unaffected | -37.4 | 6.7 | 13 |
|  | HI Mild | -38.7 | 8.1 | 9 |
|  | HI Severe | -38.1 | 10.6 | 8 |
| PIC max (mV) | Sham | -30.0 | 7.3 | 9 |
|  | HI Unaffected | -22.9 | 7.6 | 13 |
|  | HI Mild | -22.6 | 7.4 | 9 |
|  | HI Severe | -23.1 | 9.2 | 8 |
| PIC negative<br>slope range<br>(mV) | Sham | 12.9 | 3.1 | 9 |
|  | HI Unaffected | 14.4 | 2.7 | 13 |
|  | HI Mild | 16.0 | 2.8 | 9 |
|  | HI Severe | 15.0 | 3.1 | 8 |
| PIC amplitude<br>(pA) | Sham | -304 | 200 | 9 |
|  | HI Unaffected | -359 | 188 | 13 |
|  | HI Mild | -417 | 273 | 9 |
|  | HI Severe | -344 | 114 | 8 |

**Table 6.**
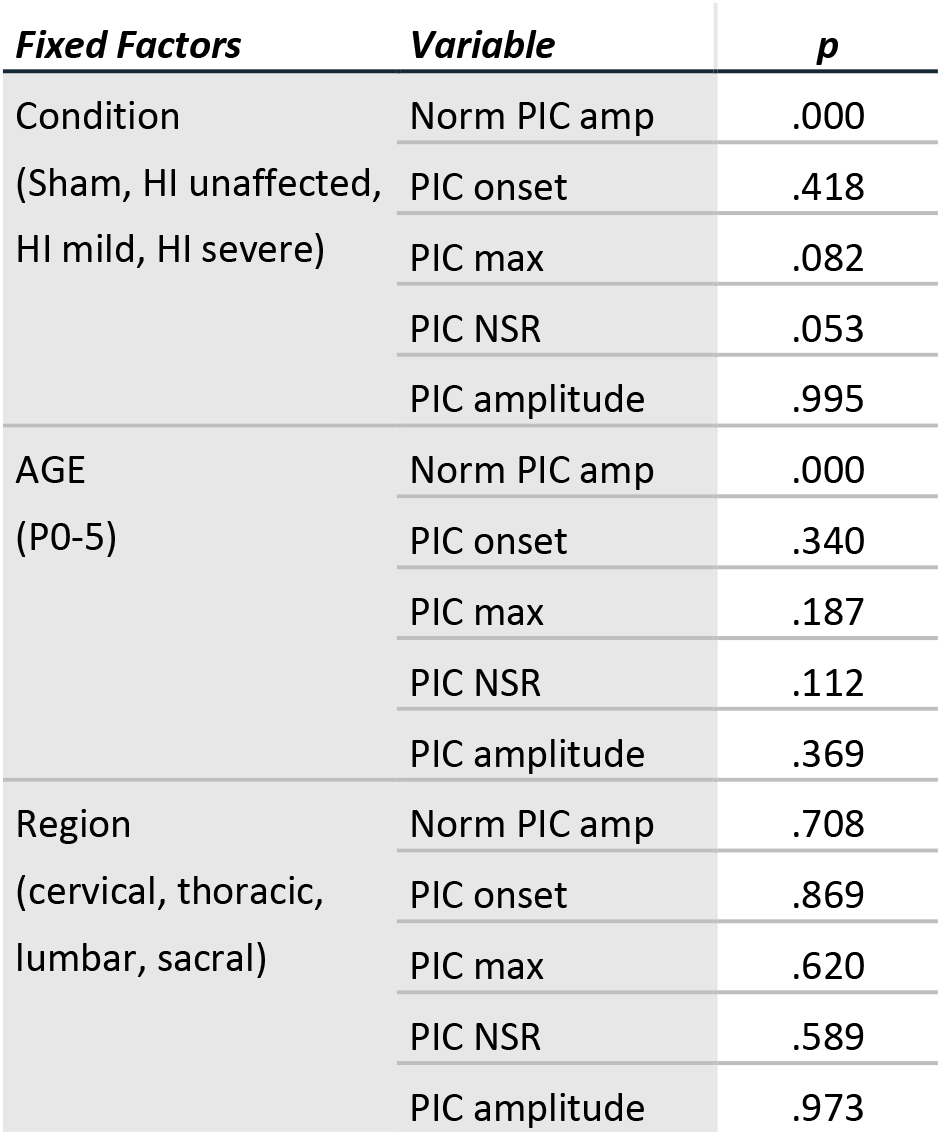
Significance of persistent inward current variables (5s ramp)

| **Fixed Factors** | **Variable** | **p** |
| --- | --- | --- |
| Condition<br>(Sham, HI unaffected,<br>HI mild, HI severe) | Norm PIC amp | .000 |
|  | PIC onset | .418 |
|  | PIC max | .082 |
|  | PIC NSR | .053 |
|  | PIC amplitude | .995 |
| AGE<br>(P0-5) | Norm PIC amp | .000 |
|  | PIC onset | .340 |
|  | PIC max | .187 |
|  | PIC NSR | .112 |
|  | PIC amplitude | .369 |
| Region<br>(cervical, thoracic,<br>lumbar, sacral) | Norm PIC amp | .708 |
|  | PIC onset | .869 |
|  | PIC max | .620 |
|  | PIC NSR | .589 |
|  | PIC amplitude | .973 |

**Table 7.** Persistent inward current characteristics (16s ramp)

| **Variable** | **Condition** | **Mean** | **SD** | **N** |
| --- | --- | --- | --- | --- |
| Norm PIC amp<br>(pA/pF) | Sham | -2.32 | 2.77 | 6 |
|  | HI Unaffected | -1.58 | .90 | 13 |
|  | HI Mild | -1.73 | .94 | 9 |
|  | HI Severe | -1.03 | .50 | 8 |
| PIC onset (mV) | Sham | -47.2 | 6.3 | 6 |
|  | HI Unaffected | -39.7 | 6.3 | 13 |
|  | HI Mild | -40.9 | 9.7 | 9 |
|  | HI Severe | -35.0 | 11.5 | 8 |
| PIC max (mV) | Sham | -27.8 | 9.9 | 6 |
|  | HI Unaffected | -21.4 | 7.4 | 13 |
|  | HI Mild | -24.2 | 9.9 | 9 |
|  | HI Severe | -18.3 | 11.1 | 8 |
| PIC negative<br>slope range<br>(mV) | Sham | 19.5 | 6.0 | 6 |
|  | HI Unaffected | 18.3 | 3.2 | 13 |
|  | HI Mild | 16.6 | 2.4 | 9 |
|  | HI Severe | 16.7 | 4.1 | 8 |
| PIC amplitude<br>(pA) | Sham | -260 | 200 | 6 |
|  | HI Unaffected | -391 | 214 | 13 |
|  | HI Mild | -374 | 253 | 9 |
|  | HI Severe | -289 | 140 | 8 |

**Table 8.** Significance of persistent inward current variables (16s ramp)

| **Fixed Factors** | **Variable** | **p** |
| --- | --- | --- |
| Condition<br>(Sham, HI unaffected,<br>HI mild, HI severe) | Norm PIC amp | .002 |
|  | PIC onset | .039 |
|  | PIC max | .033 |
|  | PIC NSR | .466 |
|  | PIC amplitude | .464 |
| AGE<br>(P0-5) | Norm PIC amp | .000 |
|  | PIC onset | .984 |
|  | PIC max | .789 |
|  | PIC NSR | .350 |
|  | PIC amplitude | .369 |
| Region<br>(cervical, thoracic,<br>lumbar, sacral) | Norm PIC amp | .897 |
|  | PIC onset | .499 |
|  | PIC max | .685 |
|  | PIC NSR | .732 |
|  | PIC amplitude | .938 |

**Table 9.**
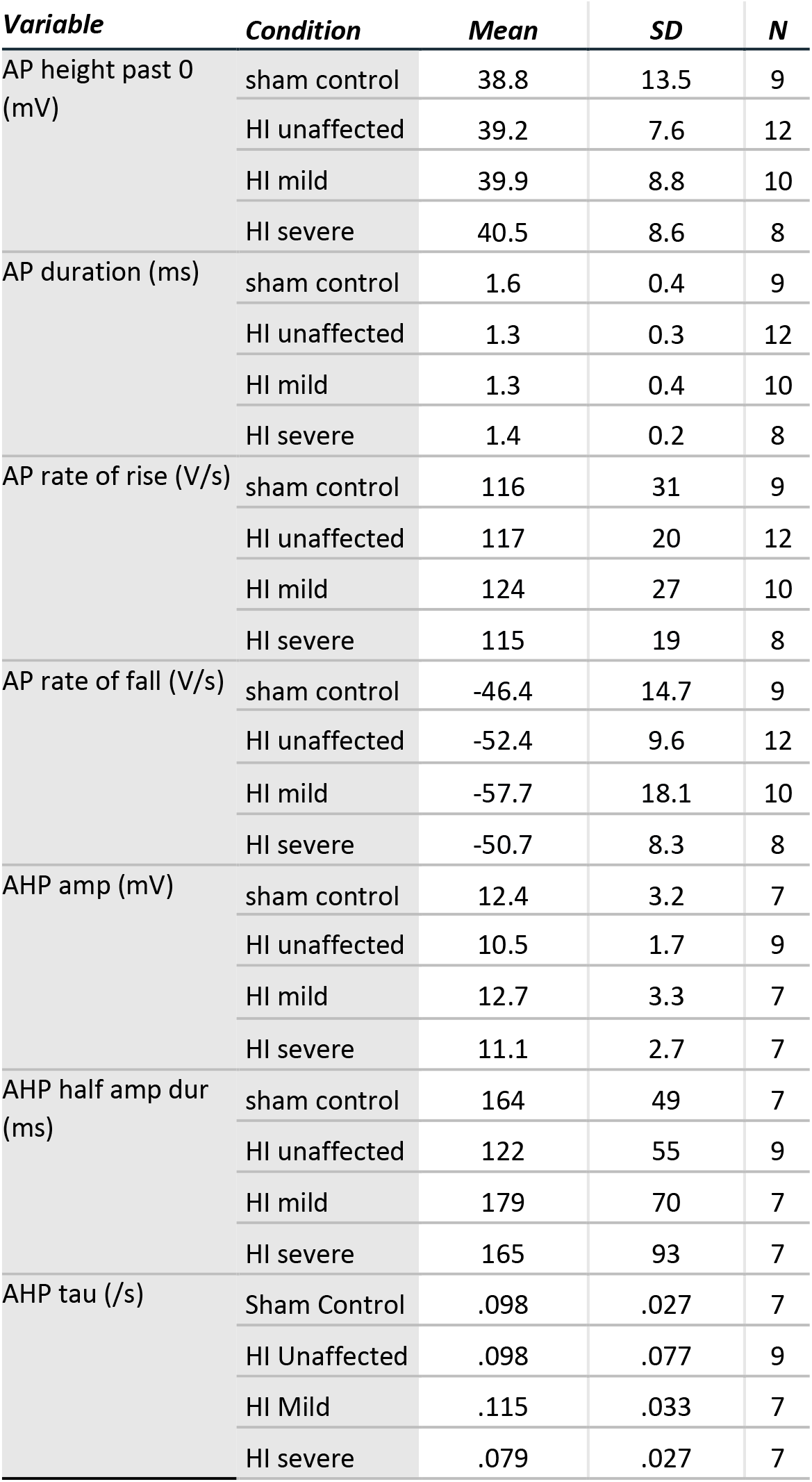
Action potential characteristics

**Table 10.** Significance of action potential variables

| **Fixed Factor** | **Variable** | **p** |
| --- | --- | --- |
| Condition<br>(Sham control, HI<br>unaffected, HI<br>mildly affected, HI<br>severely affected) | AP height past 0 | .870 |
|  | AP duration | .811 |
|  | AP rate of rise | .789 |
|  | AP rate of fall | .858 |
|  | AHP amp | .715 |
|  | AHP half amp dur | .284 |
|  | AHP tau | .437 |
| AGE<br>(P0-5) | AP height past 0 | .400 |
|  | AP duration | .192 |
|  | AP rate of rise | .267 |
|  | AP rate of fall | .118 |
|  | AHP amp | .542 |
|  | AHP half amp dur | .151 |
|  | AHP tau | .002 |
| Region<br>(cervical, thoracic,<br>lumbar, sacral) | AP height past 0 | .931 |
|  | AP duration | .312 |
|  | AP rate of rise | .435 |
|  | AP rate of fall | .287 |
|  | AHP amp | .045 |
|  | AHP half amp dur | .036 |
|  | AHP tau | .405 |

**Table 11.** Characteristics of Ih

| **Variable** | **Condition** | **Mean** | **SD** | **N** |
| --- | --- | --- | --- | --- |
| Sag (mV) | sham control | 18.2 | 15.5 | 9 |
|  | HI unaffected | 13.9 | 9.4 | 12 |
|  | HI mild | 14.2 | 7.0 | 10 |
|  | HI severe | 14.1 | 5.5 | 8 |
| Rebound (mV) | sham control | 6.1 | 5.0 | 9 |
|  | HI unaffected | 6.1 | 3.2 | 12 |
|  | HI mild | 6.6 | 2.0 | 10 |
|  | HI severe | 6.2 | 2.4 | 8 |
| Sag % | sham control | 48.4 | 32.6 | 9 |
|  | HI unaffected | 36.9 | 13.2 | 12 |
|  | HI mild | 39.6 | 20.5 | 10 |
|  | HI severe | 37.6 | 14.2 | 8 |
| Rebound % | sham control | 16.9 | 9.8 | 9 |
|  | HI unaffected | 17.2 | 6.0 | 12 |
|  | HI mild | 19.2 | 10.2 | 10 |
|  | HI severe | 17.1 | 7.5 | 8 |

**Table 12.** Significance of Ih related variables

| **Fixed Factors** | **Variable** | **p** |
| --- | --- | --- |
| Condition<br>(Sham, HI<br>unaffected, HI<br>mild, HI severe) | Sag (mV) | .816 |
|  | rebound (mV) | .665 |
|  | sag % | .934 |
|  | rebound% | .886 |
| Age<br>(P0-5) | Sag (mV) | .882 |
|  | rebound (mV) | .911 |
|  | sag % | .930 |
|  | rebound% | .924 |
| Region<br>(cervical, thoracic,<br>lumbar, sacral) | Sag (mV) | .640 |
|  | rebound (mV) | .910 |
|  | sag % | .236 |
|  | rebound% | .908 |

### Persistent inward currents suggest excitability is dampened after HI

PICs were significantly affected by hypoxia-ischemia, revealing that intrinsic excitability may be dampened. PICs were measured using both short (5s) and long (16s) protocols, which can preferentially activate and inactivate Na^+^ and Ca^2+^ mediated PICs. The different protocols yielded different results. For example, there was a non-significant trend for more depolarized voltage dependence in the PICs evoked using a short 5 second voltage ramp as shown in **Figure 3**. Using longer voltage ramps (16s), the change in voltage dependence of the PIC became significant for both PIC max and PIC onset (see Tables 7 – 8). The range of the negative slope region (PIC Max – PIC onset) also trended towards a broader voltage range in HI MNs using the 5s ramps but did not reach significance (*p* = .053). Using 16s ramps, this trend was abolished. Since changes in PIC onset and maximum voltage were more pronounced in longer ramps this could suggest an altered balance of Na^+^ and Ca^2+^ channel activation or altered activation / inactivation of these channels (see discussion). Change in the magnitude of the PIC was not observed outright in either of the protocols: the magnitude of the currents was similar between groups (see figure 3E and Tables 5 – 8), however the normalized amplitude was significantly smaller in HI MNs with both protocols (see figure 3F and Tables 5 – 8). Normalized amplitude is based on the whole cell capacitance of the MNs, which can be used as an indirect measurement of membrane area. In other words, the current generated per unit of membrane area in HI injured MNs was smaller than in control MNs. The PIC parameters for all groups are summarized in Figure 3C – F, and tables 5 – 8. Intrinsic properties of all MNs (including capacitance and input resistance) are included in Tables 3 and 4.

**Figure 3:**
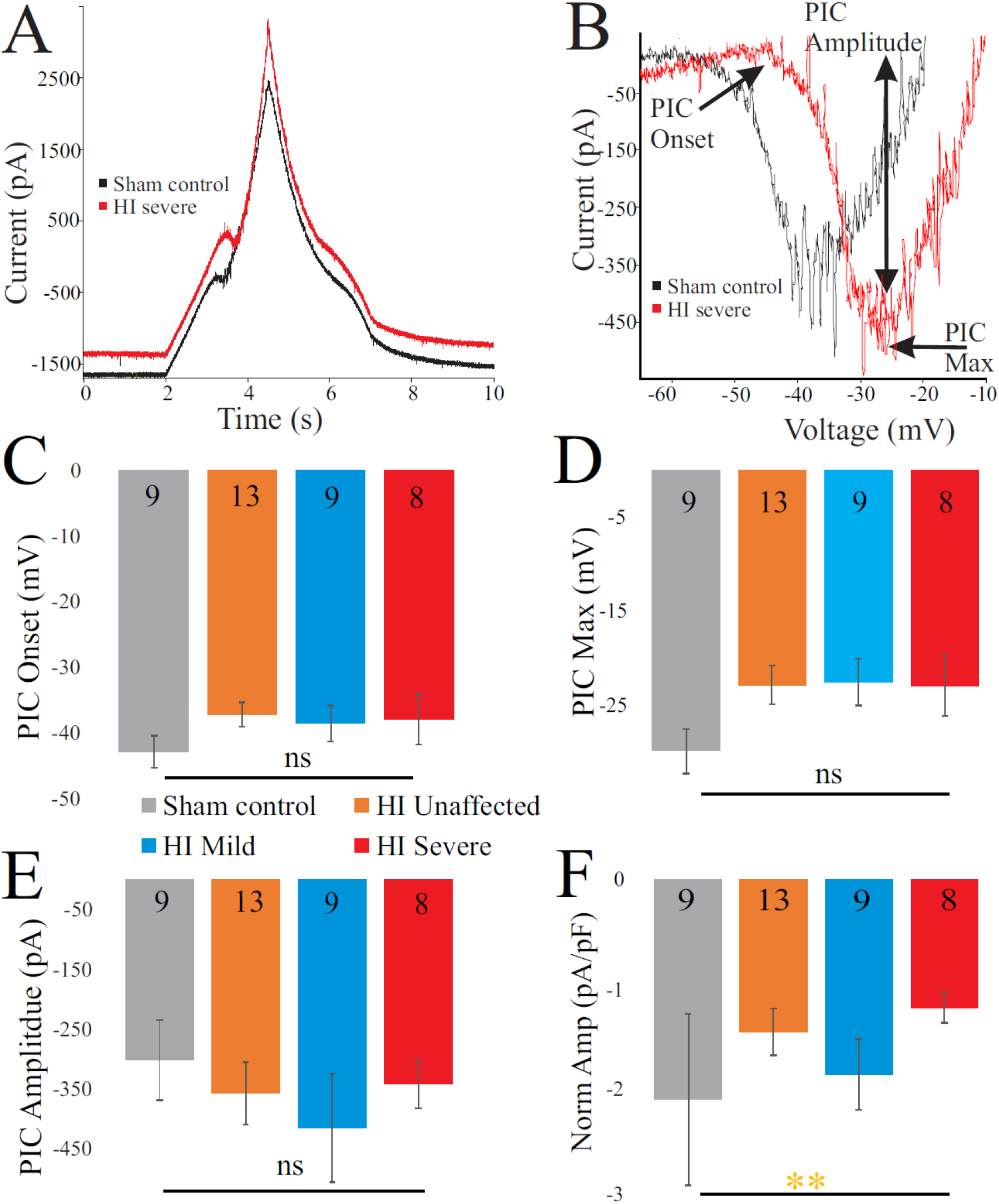
PICs are altered in HI injured motoneurons. **(A)** Typical current response to voltage ramp in a sham (black trace) and HI severe (red) MN. **(B)** Leak-subtracted PICs from sham control (black) and HI severe (red) motoneurons are similar in amplitude. There is non-significant trend for depolarized PIC onset **(C)** and **(D)** PIC Max after HI injury. **(E)** PIC Amplitude was not significantly changed. **(F)** PIC amplitude normalized to capacitance was significantly smaller after HI injury. Error bars = SEM.

### Morphology affected by HI injury

Morphology of MNs was assessed in all patched neurons, as shown in **Figure 4**. As suggested by the normalized PIC amplitude, membrane properties relating to neuron size are affected by HI injury, including a significantly larger capacitance (Fig 4C and Tables 3 and 4). These changes suggest increased neuron size after HI injury. The soma size was unchanged: there were no significant differences between groups in soma largest cross-sectional area (Fig 4D) or other measurements of soma size (Tables 13 and 14). There was, however, a significant increase in dendrite length and an increase in the number of primary dendrites in HI injured MNs, which could account for changes in electrical properties. All data pertaining to dendritic morphology is included in Tables 15 – 16.

**Figure 4:**
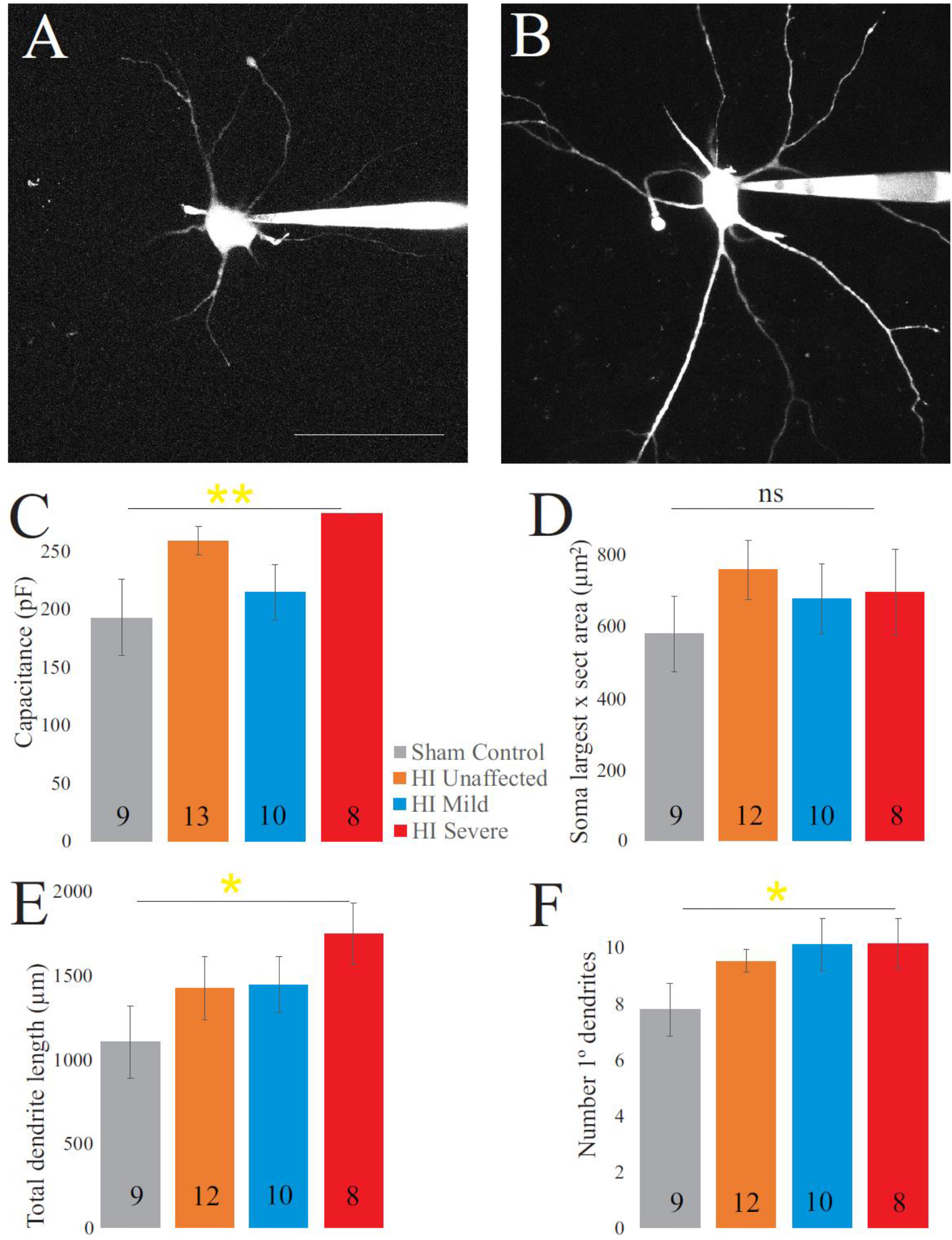
Morphology is affected by HI. Typical sham control **(A)** and HI severe **(B)** motoneurons filled with dye during patch clamp (electrodes visible on right). Average values of whole cell capacitance **(C)**, soma largest cross-sectional area **(D)**, total dendrite length **(E)**, and number of stem dendrites **(F)** are included for all neurons. N of each group is included at the base of the bar graph. Scale bar in A = 100μm, applies to A and B. Error bars = SEM.

**Table 13.**
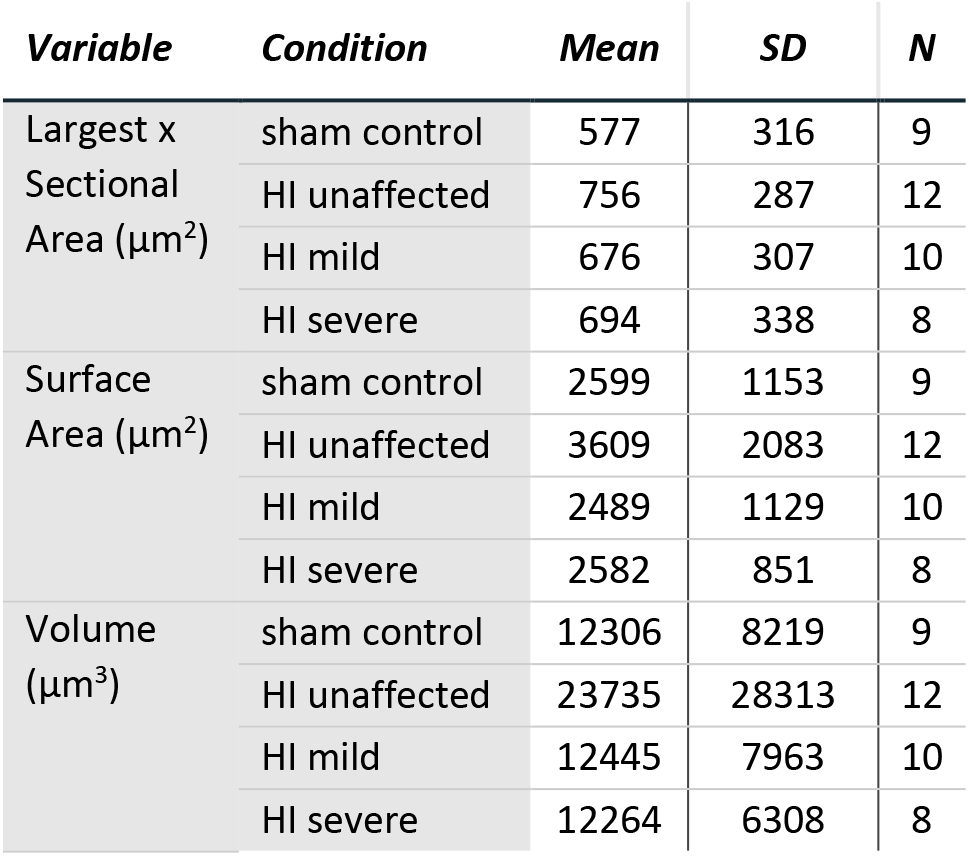
Soma morphology characteristics

| *Variable* | *Condition* | *Mean* | *SD* | *N* |
| --- | --- | --- | --- | --- |
| Largest x Sectional Area ($\mu\text{m}^2$) | sham control | 577 | 316 | 9 |
|  | HI unaffected | 756 | 287 | 12 |
|  | HI mild | 676 | 307 | 10 |
|  | HI severe | 694 | 338 | 8 |
| Surface Area ($\mu\text{m}^2$) | sham control | 2599 | 1153 | 9 |
|  | HI unaffected | 3609 | 2083 | 12 |
|  | HI mild | 2489 | 1129 | 10 |
|  | HI severe | 2582 | 851 | 8 |
| Volume ($\mu\text{m}^3$) | sham control | 12306 | 8219 | 9 |
|  | HI unaffected | 23735 | 28313 | 12 |
|  | HI mild | 12445 | 7963 | 10 |
|  | HI severe | 12264 | 6308 | 8 |

**Table 14.** Significance soma characteristics

| *Fixed Factors* | *Variable* | *p* |
| --- | --- | --- |
| Condition | Largest x Sectional Area (Soma) | .082 |
|  | Surface Area (Soma) | .212 |
|  | Volume (Soma) | .348 |
| Age | Largest x Sectional Area (Soma) | .629 |
|  | Surface Area (Soma) | .692 |
|  | Volume (Soma) | .742 |
| Region | Largest x Sectional Area (Soma) | .148 |
|  | Surface Area (Soma) | .690 |
|  | Volume (Soma) | .851 |

**Table 15.** Dendrite characteristics

| **Variable** | **Condition** | **Mean** | **SD** | **N** |
| --- | --- | --- | --- | --- |
| Number of Dendrites | sham control | 7.7 | 2.8 | 9 |
|  | HI unaffected | 9.5 | 1.4 | 12 |
|  | HI mild | 10.1 | 2.9 | 10 |
|  | HI severe | 10.1 | 2.5 | 8 |
| Nodes | sham control | 14.8 | 14.1 | 9 |
|  | HI unaffected | 14.7 | 11.5 | 12 |
|  | HI mild | 13.5 | 5.7 | 10 |
|  | HI severe | 17.0 | 8.9 | 8 |
| Length ($\mu\text{m}$) | sham control | 1103 | 649 | 9 |
|  | HI unaffected | 1422 | 664 | 12 |
|  | HI mild | 1443 | 537 | 10 |
|  | HI severe | 1744 | 521 | 8 |
| Mean Length ($\mu\text{m}$) | sham control | 140 | 67 | 9 |
|  | HI unaffected | 149 | 67 | 12 |
|  | HI mild | 144 | 32 | 10 |
|  | HI severe | 178 | 57 | 8 |
| Surface Area ($\mu\text{m}^2$) | sham control | 6264 | 3842 | 9 |
|  | HI unaffected | 9667 | 7000 | 12 |
|  | HI mild | 7537 | 3293 | 10 |
|  | HI severe | 9868 | 5384 | 8 |
| Mean Surface ($\mu\text{m}^2$) | sham control | 792 | 414 | 9 |
|  | HI unaffected | 1024 | 772 | 12 |
|  | HI mild | 740 | 159 | 10 |
|  | HI severe | 1012 | 603 | 8 |
| Volume ($\mu\text{m}^3$) | sham control | 3614 | 2642 | 9 |
|  | HI unaffected | 6926 | 7512 | 12 |
|  | HI mild | 4073 | 2406 | 10 |
|  | HI severe | 5844 | 5239 | 8 |
| Mean Volume ($\mu\text{m}^3$) | sham control | 451 | 295 | 9 |
|  | HI unaffected | 743 | 836 | 12 |
|  | HI mild | 390 | 129 | 10 |
|  | HI severe | 605 | 587 | 8 |

**Table 16.** Significance dendrite characteristics

| **Fixed Factor** | **Variable** | **p** |
| --- | --- | --- |
| Condition<br>(Sham, HI<br>unaffected, HI<br>mild, HI severe) | # Dendrites | .046 |
|  | Nodes | .081 |
|  | Length (Den) | .021 |
|  | Mean Length (Den) | .312 |
|  | Surface Area (Den) | .216 |
|  | Mean Sur (Den) | .580 |
|  | Volume (Den) | .461 |
|  | Mean Volume (Den) | .643 |
| AGE<br>(P0-5) | # Dendrites | .687 |
|  | Nodes | .163 |
|  | Length (Den) | .698 |
|  | Mean Length (Den) | .561 |
|  | Surface Area (Den) | .643 |
|  | Mean Sur (Den) | .563 |
|  | Volume (Den) | .695 |
|  | Mean Volume (Den) | .659 |
| Region<br>(cervical,<br>thoracic,<br>lumbar, sacral) | # Dendrites | .700 |
|  | Nodes | .009 |
|  | Length (Den) | .228 |
|  | Mean Length (Den) | .062 |
|  | Surface Area (Den) | .532 |
|  | Mean Sur (Den) | .352 |
|  | Volume (Den) | .756 |
|  | Mean Volume (Den) | .659 |

## Discussion

### Summary

Electrophysiological properties of spinal MNs are altered by developmental HI injury, and the magnitude of changes are correlated to severity of motor deficits. Specifically, these changes include increased sustained firing and a higher threshold for action potentials. These differences could contribute to both problems of weakness and muscle stiffness that is common in spastic cerebral palsy. Since traditional views of CP largely view motor dysfunction as a result of the damaged motor cortex improperly signaling to spinal neurons, our new evidence suggests this is only part of the problem. Spinal MNs are not developing the same after HI and show an overall change in excitability. Whether these changes are directly due to the HI insult or indirectly due to downstream effects must be determined by future work.

### Contribution of spinal motoneurons to dysfunction in cerebral palsy

Here we show that MNs are more excitable after HI injury, including elevated resting potential, and more sustained firing. Previous work showed that after HI injury in rabbits there were also fewer spinal MNs, and spinal interneurons in lamina VII were undergoing apoptosis (15). After loss of corticospinal projections, it was recently found that the spinal cholinergic interneurons which give rise to C boutons on MNs are lost (33–35). Taken together this data suggests spinal circuits are 1) just as vulnerable to HI injury as the developing cortex and 2) potentially functioning with fewer neurons and altered circuitry. In addition to fewer neurons, there is also atrophy of the muscles which could contribute to motor deficits in cerebral palsy. In mice, rabbits, and humans, muscle atrophy appears along with losses in numbers of MNs (7, 15, 28, 42). Recent work has shown similar changes to muscle architecture in the rabbit HI model of CP to humans, including atrophy and shortening along with longer sarcomere length. Increased muscle stiffness in rabbits affected by HI was found even after administration of anesthetic – indicating some muscle stiffness is derived from mechanical changes in the muscles, though a large component of the muscle stiffness was diminished with anesthetic thus was driven neurally (54). During development both feedback and feed-forward signaling can regulate growth and maturation, processes which may be disrupted in CP in both MNs and muscle fibers. It is likely the loss of spinal interneurons, MNs and muscle fibers reduces coordination and strength in those with cerebral palsy. Altered size and excitability of MNs is a feature of other motor disorders including amyotrophic lateral sclerosis and spinal muscular atrophy (17, 24, 49, 50, 53). This data supports further exploration of interventions for CP and other motor disorders that target spinal MNs, neuromodulators which could alter spinal circuits, and therapies aimed at restoring balance within spinal circuits for alleviation of spasticity.

### Neuromodulation

The exact causes of the changes in MN physiology observed here are unclear, but they could result from the increase in spinal monoamines that occurs after developmental HI injury in both rodents and rabbits (3, 16). Serotonin is generally thought of as a neurotransmitter and neuromodulator, but developmental disruption in 5HT is associated with neurological disorders including autism, Rett syndrome, Down’s syndrome and, more recently, cerebral palsy (1, 3, 13, 16, 45, 57, 59, 61). Serotonin increases MN excitability in neonatal and juvenile mice, rats and guinea pigs (30, 31, 56, 62), so it likely has the same effect on rabbit MNs. Depolarization of the resting membrane potential, increased action potential firing through hyperpolarization of the voltage threshold and enhanced PIC, increased action potential height and reduction of high-voltage activated Ca^2+^ entry are all associated with 5HT receptor activation in neonatal and adult MNs (2, 18, 23, 30–32, 36, 38, 39). Therefore increased 5HT could have a direct impact on excitability, though in HI rabbits the increase in 5HT was accompanied by decreased mRNA for 5HT2 receptors and increased mRNA for the SERT serotonin transporter (16). In light of that finding, it is not clear that neurons remain responsive to 5HT. In the experiments here, all MNs were recorded in spinal cord slices incubated and perfused in standard oxygenated aCSF without any serotonergic drugs present. Therefore, HI MNs *in vivo* could show different levels of excitability since they would be in the presence of elevated 5HT, while our MNs were all recorded in the same physiological solution. Thus the contribution of 5HT to the altered excitability observed here is restricted to its chronic effects on neuron development, namely morphological changes. Serotonin 5HT1A and 5HT2A receptor activation increases neurite outgrowth, dendritic branching, and spine formation (6, 19, 44), findings that align well with the present finding of increased dendritic length and number of primary dendrites in the HI MNs. Future experiments will be needed to address the role of 5HT in enhancing MN excitability and its effects on synaptically-evoked action potentials. Synaptic events in dendrites would more strongly evoke PICs, though both altered dendritic morphology and elevated 5HT could dampen them.

### Possible mechanism of altered MN output

The mechanism for increased MN activity and thus muscle stiffness may be due to delayed Na^+^ channel inactivation, or an increased contribution of Ca^2+^ to the PICs in severe HI MNs. In neonatal MNs, Na^+^ channels generate the majority of the PIC and account for action potential initiation / ability to repetitively fire. Typically Na^+^ channels contributing to the PIC and repetitive firing (Nav 1.1, 1.2 and 1.6) (4, 5, 52) inactivate faster than the corresponding Ca^2+^ channels (38, 48). Short voltage ramps preferentially measure the Na^+^ PIC for this reason: Na^+^ channels inactivate quickly enough that even on the descending ramp of the short protocol, there is no longer a region of negative slope (see figure 3A). Changes in Na^+^ channel inactivation in adult MNs along with postnatal development of the longer-lasting Ca^2+^ PIC makes typical adult MNs display more negative ΔI values and longer lasting PICs (29, 38, 50). In the neonatal rabbit MNs, sham controls showed positive ΔI values that are quite typical for neonates, while HI MNs showed significantly more negative values. This could be due to slower Na^+^ channel inactivation or increased contribution of Ca^2+^ channels to the PICs after injury. To fully determine the altered mechanism of aberrant firing after HI injury future studies into the biophysical properties of Na^+^ channels and maturation of Ca^2+^ channel expression must be pursued.

### Severity in motor deficits and electrophysiology

Generally, unaffected and mildly affected MNs showed parameters that were intermediate between control MNs and severe MNs. However, in one category, the frequency-current relationship, HI unaffected MNs showed the most prominent changes. Thus electrophysiological changes were overwhelmingly in line with phenotype, suggesting abberrent MN properties could be contributing to the severity of the phenotype. It cannot be ruled out, however, that HI “unaffected” MNs may have a subtle phenotype that is not readily evident based on the behavioral testing we performed here but can be detected with patch clamp recording. Or perhaps abnormalities in these rabbits would develop in later in life: in humans CP patients, diagnosis of CP is not made until 18-24 months of life and the peak of spasticity occurs around four years of age (26, 27, 47). Future work is needed to assess maturation of the MN properties in different groups, the potential contribution of delayed Na^+^ channel inactivation in CP, progression of motor deficits with age, and the development of new therapeutic strategies that could target MNs.

### Conclusion

Changes in MN physiology after developmental injury are consistent with motor deficits in rabbits. This suggests not only brain injuries but also changes in the spinal cord contribute to impaired function in cerebral palsy. Exploring both altered maturation of spinal neurons and loss of descending connectivity should be pursued to improve outcomes for individuals with cerebral palsy.

## References

1. Bar-Peled O, Gross-Isseroff R, Ben-Hur H, Hoskins I, Groner Y, Biegon A. Fetal human brain exhibits a prenatal peak in the density of serotonin 5-HT1A receptors. Neurosci Lett 127: 173–176, 1991.

2. Bayliss DA, Umemiya M, Berger AJ. Inhibition of N- and P-type calcium currents and the after-hyperpolarization in rat motoneurones by serotonin. J Physiol 485: 635–647, 1995.

3. Bellot B, Peyronnet-Roux J, Gire C, Simeoni U, Vinay L, Viemari JC. Deficits of brainstem and spinal cord functions after neonatal hypoxia-ischemia in mice. Pediatr Res 75: 723–730, 2014.

4. Boiko T, Rasband MN, Rock S, Caldwell JH, Mandel G, Trimmer JS, Matthews G, Brook S, York N, Caldwell R. Compact Myelin Dictates the Differential Targeting of Two Sodium Channel Isoforms in the Same Axon The State University of New York at Stony Brook. 30: 91–104, 2001.

5. Boiko T, Wart A Van, Caldwell JH, Levinson SR, Trimmer JS, Matthews G. Functional Specialization of the Axon Initial Segment by Isoform-Specific Sodium Channel Targeting. 23: 2306–2313, 2003.

6. Bou-Flores C, Lajard A, Monteau R, De Maeyer E, Seif I, Lanoir J, Hilaire G. Abnormal Phrenic Motoneuron Activity and Morphology in Neonatal Monoamine Oxidase A-Deficient Transgenic Mice: Possible Role of a Serotonin Excess. 20: 4646–4656, 2000.

7. Brandenburg XJE, Gransee HM, Fogarty MJ, Sieck GC. Differences in lumbar motor neuron pruning in an animal model of early onset spasticity. (2019). doi: 10.1152/jn.00186.2018.

8. Buser JR, Segovia KN, Dean JM, Nelson K, Beardsley D, Gong X, Luo NL, Ren J, Wan Y, Riddle A, McClure MM, Ji X, Derrick M, Hohimer AR, Back SA, Tan S. Timing of appearance of late oligodendrocyte progenitors coincides with enhanced susceptibility of preterm rabbit cerebral white matter to hypoxia-ischemia. J Cereb Blood Flow Metab 30: 1053–1065, 2010.

9. Cavarsan CF, Gorassini MA, Quinlan KA. Animal models of developmental motor disorders: parallels to human motor dysfunction in cerebral palsy. J Neurophysiol 122: 1238–1253, 2019.

10. Clowry GJ. The dependence of spinal cord development on corticospinal input and its significance in understanding and treating spastic cerebral palsy. Neurosci Biobehav Rev 31: 1114–1124, 2007.

11. Clowry GJ, Basuodan R, Chan F. What are the best animal models for testing early intervention in cerebral palsy? Front Neurol 5: 1–17, 2014.

12. Clowry GJ, Davies BM, Upile NS, Gibson CL, Bradley PM. Spinal cord plasticity in response to unilateral inhibition of the rat motor cortex during development: Changes to gene expression, muscle afferents and the ipsilateral corticospinal projection. Eur J Neurosci 20: 2555–2566, 2004.

13. De Filippis B, Chiodi V, Adriani W, Lacivita E, Mallozzi C, Leopoldo M, Domenici MR, Fuso A, Laviola G. Long-lasting beneficial effects of central serotonin receptor 7 stimulation in female mice modeling Rett syndrome. Front Behav Neurosci 9: 1–11, 2015.

14. Derrick M, Luo NL, Bregman JC, Jilling T, Ji X, Fisher K, Gladson CL, Beardsley DJ, Murdoch G, Back SA, Tan S. Preterm Fetal Hypoxia-Ischemia Causes Hypertonia and Motor Deficits in the Neonatal Rabbit: A Model for Human Cerebral Palsy? J Neurosci 24: 24–34, 2004.

15. Drobyshevsky A, Quinlan KA. Spinal cord injury in hypertonic newborns after antenatal hypoxia-ischemia in a rabbit model of cerebral palsy. Exp Neurol 293: 13–26, 2017.

16. Drobyshevsky A, Takada SH, Luo K, Derrick M, Yu L, Quinlan KA, Vasquez-Vivar J, Nogueira MI, Tan S. Elevated spinal monoamine neurotransmitters after antenatal hypoxia-ischemia in rabbit cerebral palsy model. J Neurochem 132: 394–402, 2015.

17. Dukkipati SS, Garrett TL, Elbasiouny SM. The vulnerability of spinal motoneurons and soma size plasticity in a mouse model of amyotrophic lateral sclerosis. J Physiol 596: 1723–1745, 2018.

18. Elliot P, Wallis DI. Serotonin and L-Norepinephrine as Mediators of Altered Excitability in Neonatal Rat Motoneurons Studied In Vitro. Neuroscience 47: 533–544, 1992.

19. Fricker AD, Rios C, Devi LA, Gomes I. Serotonin receptor activation leads to neurite outgrowth and neuronal survival. Mol Brain Res 138: 228–235, 2005.

20. Friel K, Chakrabarty S, Kuo HC, Martin J. Using motor behavior during an early critical period to restore skilled limb movement after damage to the corticospinal system during development. J Neurosci 32: 9265–9276, 2012.

21. Friel KM, Martin JH. Role of sensory-motor cortex activity in postnatal development of corticospinal axon terminals in the cat. J Comp Neurol 485: 43–56, 2005.

22. Friel KM, Martin JH. Bilateral activity-dependent interactions in the developing corticospinal system. J Neurosci 27: 11083–11090, 2007.

23. Gilmore J, Fedirchuk B. The excitability of lumbar motoneurones in the neonatal rat is increased by a hyperpolarization of their voltage threshold for activation by descending serotonergic fibres. J Physiol 558: 213–224, 2004.

24. Gogliotti RG, Quinlan KA, Barlow CB, Heier CR, Heckman CJ, DiDonato CJ. Motor neuron rescue in spinal muscular atrophy mice demonstrates that sensory-motor defects are a consequence, not a cause, of motor neuron dysfunction. J Neurosci 32: 3818–3829, 2012.

25. Graham HK, Rosenbaum P, Paneth N, Dan B, Lin J. Graham et al. - 2016 - Cerebral palsy. (2016). doi: 10.1038/nrdp.2015.82.

26. Hadders-Algra M. Early diagnosis and early intervention in cerebral palsy. Front Neurol 5: 1–13, 2014.

27. Hägglund G, Wagner P. Development of spasticity with age in a total population of children with cerebral palsy. BMC Musculoskelet Disord 9: 1–10, 2008.

28. Han Q, Feng J, Qu Y, Ding Y, Wang M, So KF, Wu W, Zhou L. Spinal cord maturation and locomotion in mice with an isolated cortex. Neuroscience 253: 235–244, 2013.

29. Harvey PJ, Li X, Li Y, Bennett DJ. 5-HT 2 receptor activation facilitates a persistent sodium current and repetitive firing in spinal motoneurons of rats with and without chronic spinal cord injury. J Neurophysiol 96: 1158–1170, 2006.

30. Hsiao CF, Del Negro CA, Trueblood PR, Chandler SH. Ionic basis for serotonin-induced bistable membrane properties in guinea pig trigeminal motoneurons. J Neurophysiol 79: 2847–2855, 1998.

31. Hsiao CF, Trueblood PR, Levine MS, Chandler SH. Multiple effects of serotonin on membrane properties of trigeminal motoneurons in vitro. J Neurophysiol 77: 2910–2924, 1997.

32. Inoue T, Itoh S, Kobayashi M, Kang Y, Matsuo R, Wakisaka S, Morimoto T. Serotonergic modulation of the hyperpolarizing spike afterpotential in rat jaw-closing motoneurons by PKA and PKC. J Neurophysiol 82: 626–637, 1999.

33. Jiang Y-Q, Armada K, Martin JH. Neuronal activity and microglial activation support corticospinal tract and proprioceptive afferent sprouting in spinal circuits after a corticospinal system lesion. Exp Neurol 321: 113015, 2019.

34. Jiang YQ, Sarkar A, Amer A, Martin JH. Transneuronal downregulation of the premotor cholinergic system after corticospinal tract loss. J Neurosci 38: 8329–8344, 2018.

35. Jiang YQ, Zaaimi B, Martin JH. Competition with primary sensory afferents drives remodeling of corticospinal axons in mature spinal motor circuits. J Neurosci 36: 193–203, 2016.

36. Larkman PM, Kelly JS. Ionic Mechanisms Mediating 5-Hydroxytryptamine- and Noradrenaline-Evoked Depolarization of Adult Rat Facial Motoneurones. J Physiol 456: 473–490, 1992.

37. Li Q, Martin JH. Postnatal development of differential projections from the caudal and rostral motor cortex subregions. Exp Brain Res 134: 187–198, 2000.

38. Li X, Murray K, Harvey PJ, Ballou EW, Bennett DJ. Serotonin facilitates a persistent calcium current in motoneurons of rats with and without chronic spinal cord injury. J Neurophysiol 97: 1236–1246, 2007.

39. Lindsay AD, Feldman JL. Modulation of respiratory activity of neonatal rat phrenic motoneurones by serotonin. J Physiol 461: 213–233, 1993.

40. MacLennon A, International Cerebral Palsy Task Force. A template for definint a causal relation between acute intrapartum events and cerebral palsy: international consensus statement. Br Med J 319: 1054–1059, 1999.

41. Magee L, Sawchuck D, Synnes A, van Dadelszen P, Committee TMS for FNC, Committee SMFM. Magnesium Sulphate for Fetal Neuroprotection. J Obstet Gynaecol Canada 33: 516–529, 2011.

42. Marciniak C, Li X, Zhou P. An examination of motor unit number index in adults with cerebral palsy. J Electromyogr Kinesiol 25: 444–450, 2015.

43. Martin JH, Kably B, Hacking A. Activity-dependent development of cortical axon terminations in the spinal cord and brain stem. Exp Brain Res 125: 184–199, 1999.

44. Mogha A, Guariglia SR, Debata PR, Wen GY, Banerjee P. Serotonin 1A receptor-mediated signaling through ERK and PKCα is essential for normal synaptogenesis in neonatal mouse hippocampus. Transl Psychiatry 2: 1–12, 2012.

45. Muller CL, Anacker AMJ, Veenstra-VanderWeele J. The serotonin system in autism spectrum disorder: From biomarker to animal models. Neuroscience 321: 24–41, 2016.

46. Novak I, Mcintyre S, Morgan C, Campbell L, Dark L, Morton N, Stumbles E, Wilson SA, Goldsmith S. A systematic review of interventions for children with cerebral palsy: State of the evidence. Dev Med Child Neurol 55: 885–910, 2013.

47. Novak I, Morgan C, Adde L, Blackman J, Boyd RN, Brunstrom-Hernandez J, Cioni G, Damiano D, Darrah J, Eliasson AC, De Vries LS, Einspieler C, Fahey M, Fehlings D, Ferriero DM, Fetters L, Fiori S, Forssberg H, Gordon AM, Greaves S, Guzzetta A, Hadders-Algra M, Harbourne R, Kakooza-Mwesige A, Karlsson P, Krumlinde-Sundholm L, Latal B, Loughran-Fowlds A, Maitre N, McIntyre S, Noritz G, Pennington L, Romeo DM, Shepherd R, Spittle AJ, Thornton M, Valentine J, Walker K, White R, Badawi N. Early, accurate diagnosis and early intervention in cerebral palsy: Advances in diagnosis and treatment. JAMA Pediatr 171: 897–907, 2017.

48. Perrier J françois, Hounsgaard J. 5-HT2 receptors promote plateau potentials in turtle spinal motoneurons by facilitating an L-type calcium current. J Neurophysiol 89: 954–959, 2003.

49. Quinlan KA, Reedich E, Arnold WD, Puritz A, Cavarsan CF, Heckman C, DiDonato CJ. Hyperexcitability precedes motoneuron loss in the Smn 2B/-mouse model of spinal muscular atrophy. J Neurophysiol 122: 1297–1311, 2019.

50. Quinlan KA, Schuster JE, Fu R, Siddique T, Heckman CJ. Altered postnatal maturation of electrical properties in spinal motoneurons in a mouse model of amyotrophic lateral sclerosis. J Physiol 589: 2245–2260, 2011.

51. Rouse DJ, Gibbins KJ. Magnesium sulfate for cerebral palsy prevention. Semin Perinatol 37: 414–416, 2013.

52. Rush AM, Dib-Hajj SD, Waxman SG. Electrophysiological properties of two axonal sodium channels, Nav1.2 and Nav1.6, expressed in mouse spinal sensory neurones. J Physiol 564: 803–815, 2005.

53. Shoenfeld L, Westenbroek RE, Fisher E, Quinlan KA, Tysseling VM, Powers RK, Heckman CJ, Binder MD. Soma size and Cav1.3 channel expression in vulnerable and resistant motoneuron populations of the SOD1G93A mouse model of ALS. Physiol Rep 2: 1–13, 2014.

54. Synowiec S, Lu J, Yu L, Goussakov I, Lieber R, Drobyshevsky A. Spinal hyper-excitability and altered muscle structure contribute to muscle hypertonia in newborns after antenatal hypoxia-ischemia in a rabbit cerebral palsy model. Front Neurol 10: 1–18, 2019.

55. Thoresen M, Bågenholm R, Løberg EM, Apricena F, Kjellmer I. Posthypoxic cooling of neonatal rats provides protection against brain injury. Arch Dis Child 74: 0–5, 1996.

56. Wang M, Dun N. 5-Hyroxytryptamine responses in neonate rat motoneurones in vitro. J Physiol 430: 87–103, 1990.

57. Whittle N, Sartori SB, Dierssen M, Lubec G, Singewald N. Fetal Down syndrome brains exhibit aberrant levels of neurotransmitters critical for normal brain development. Pediatrics 120: e1465–e1471, 2007.

58. Wimalasundera N, Stevenson VL. Cerebral palsy. Pract Neurol 16: 184–194, 2016.

59. Wirth A, Holst K, Ponimaskin E. How serotonin receptors regulate morphogenic signalling in neurons. Prog Neurobiol 151: 35–56, 2017.

60. Yager J, Towfighi J, Vannucci RC. Influence of mild hypothermia on hypoxic-ischemic brain damage in the immature rat. Pediatr Res 34: 525–529, 1993.

61. Yang CJ, Tan HP, Du YJ. The developmental disruptions of serotonin signaling may involved in autism during early brain development. Neuroscience 267: 1–10, 2014.

62. Ziskind-Conhaim L, Seebach BS, Gao BX. Changes in serotonin-induced potentials during spinal cord development. J Neurophysiol 69: 1338–1349, 1993.

